# Limited accessibility of nitrogen supplied as amino acids, amides, and amines as energy sources for marine *Thaumarchaeota*

**DOI:** 10.1101/2021.07.22.453390

**Authors:** Julian Damashek, Barbara Bayer, Gerhard J. Herndl, Natalie J. Wallsgrove, Tamara Allen, Brian N. Popp, James T. Hollibaugh

**Author notes:** **Corresponding authors:** Julian Damashek, 1600 Burrstone Road, Utica, NY 13502 •, •, James T. Hollibaugh, 325 Sanford Drive, Athens, GA 30602 •, •.

## Abstract

Genomic and physiological evidence from some strains of ammonia-oxidizing *Thaumarchaeota* demonstrate their additional ability to oxidize nitrogen (N) supplied as urea or cyanate, fueling conjecture about their ability to conserve energy by directly oxidizing reduced N from other dissolved organic nitrogen (DON) compounds. Similarly, field studies have shown rapid oxidation of polyamine-N in the ocean, but it is unclear whether *Thaumarchaeota* oxidize polyamine-N directly or whether heterotrophic DON remineralization is required. We tested growth of two marine *Nitrosopumilus* isolates on DON compounds including polyamines, amino acids, primary amines, and amides as their sole energy source. Though axenic cultures only consumed N supplied as ammonium or urea, there was rapid but inconsistent oxidation of N from the polyamine putrescine when cultures included a heterotrophic bacterium. Surprisingly, axenic cultures oxidized ^15^N-putrescine during growth on ammonia, suggesting co-metabolism or accelerated breakdown of putrescine by reactive metabolic byproducts. Nitric oxide, hydrogen peroxide, or peroxynitrite did not oxidize putrescine in sterile seawater. These data suggest that the N in common DON molecules is not directly accessible to marine *Thaumarchaeota*, with thaumarchaeal oxidation (and presumably assimilation) of DON-N requiring initial heterotrophic remineralization. However, reactive byproducts or enzymatic co-metabolism may facilitate limited thaumarchaeal DON-N oxidation.

## INTRODUCTION

While chemoautotrophy supported by ammonia (NH_3_) oxidation is the primary metabolism of most *Thaumarchaeota* (Francis *et al*., 2007; Stahl and de la Torre, 2012; Santoro *et al*., 2019), there has also been speculation on their additional capacity to oxidize nitrogen (N) from a variety of dissolved organic nitrogen (DON) compounds. Numerous experiments have demonstrated that some *Thaumarchaeota* can grow via oxidation of N supplied as urea: following the discovery of urease and urea transporter genes in the sponge symbiont *Cenarchaeum symbiosum* (Hallam *et al*., 2006), thaumarchaeal urease genes were documented throughout the ocean (Yakimov *et al*., 2011; Alonso-Saez *et al*., 2012; Smith *et al*., 2016), rate measurements using ^15^N tracers showed moderate urea-N oxidation in coastal waters (Tolar *et al*., 2017; Damashek *et al*., 2019a; Kitzinger *et al*., 2019; Laperriere *et al*., 2021), and marine *Thaumarchaeota* capable of growth via stoichiometric oxidation of urea-N to nitrite (NO_2_^−^) were isolated (Qin *et al*., 2014; Bayer *et al*., 2016; Carini *et al*., 2018). In addition to urea, a thaumarchaeote isolated from a hot spring contains a cyanate hydratase gene (*cynS*) and can grow via oxidation of cyanate-N (Palatinszky *et al*., 2015). Oxidation of both urea- and cyanate-N by *Thaumarchaeota* appears to drive NO_2_^−^ production in the northern Gulf of Mexico (Kitzinger *et al*., 2019). These studies provide ample evidence of widespread thaumarchaeal oxidation of urea-N, and likely cyanate-N, throughout the ocean.

In addition to urea and cyanate, there is interest in whether *Thaumarchaeota* can directly oxidize N from other common DON compounds. Early studies of marine microbial assemblages demonstrated thaumarchaeal amino acid assimilation (Ouverney and Fuhrman, 2000; Teira *et al*., 2006; Kirchman *et al*., 2007), and recent experiments in the coastal Pacific Ocean demonstrated assimilation of N but not carbon (C) from amino acids, prompting speculation about the ability of *Thaumarchaeota* to oxidize amino acid-N directly (Dekas *et al*., 2019). However, data on thaumarchaeal DON-N oxidation rates are scarce. ^15^N-tracer experiments in the coastal South Atlantic Bight, where *Thaumarchaeota* are the dominant ammonia oxidizers (Hollibaugh *et al*., 2011; Hollibaugh *et al*. 2014; Tolar *et al*., 2017; Liu *et al*., 2018; Damashek *et al*., 2019a), not only measured detectable oxidation of N supplied as urea and L-glutamic acid (L-GLU), but found oxidation rates of N from the polyamine putrescine (1,4-diaminobutane; PUT) to be far higher than oxidation rates of urea- or amino acid-N (Damashek *et al*., 2019a). Polyamines consist of a reduced C backbone with at least two amine substitutions (Tabor and Tabor, 1984). Ubiquitously high intracellular concentrations (∼mM) of polyamines such as PUT, spermine, and spermidine (Tabor and Tabor, 1985; Liu *et al*., 2016) and rapid turnover rates in seawater (Lee and Jørgensen, 1995; Liu *et al*., 2015) suggest fast microbial cycling of polyamines in the marine environment, including oxidation of polyamine-N (Damashek *et al*., 2019a). However, it is unknown whether *Thaumarchaeota* can conserve energy by oxidizing polyamine- or amino acid-N directly, or if prior heterotrophic remineralization to ammonium (NH_4_^+^) is required.

We tested the ability of two thaumarchaeal strains originally isolated from the northern Adriatic Sea (*Nitrosopumilus piranensis* D3C and *N. adriaticus* NF5; Bayer *et al*., 2016; Bayer *et al*., 2019a) to grow using a variety of DON compounds as sole energy sources. Given prior evidence of rapid PUT-N oxidation in the field, our primary focus was polyamines. Axenic thaumarchaeal cultures were unable to grow when supplied with single DON compounds, other than the expected growth on urea by *N. piranensis* D3C. However, experiments with ^15^N-labeled compounds demonstrated that axenic cultures of both strains oxidized a significant amount of PUT-N when growing on NH_3_, suggesting PUT may be co-metabolized or broken down by reactive byproducts produced during thaumarchaeal growth. Furthermore, enrichment cultures containing a heterotrophic bacterium occasionally showed rapid PUT-N oxidation, indicating an important role for heterotrophic DON remineralization in this process. Our data indicate that oxidation of reduced N by marine *Nitrosopumilaceae* is restricted to well-known substrates, but suggest breakdown of DON and oxidation of its liberated N may occur as an indirect effect of thaumarchaeal growth or due to tight coupling between DON remineralization by heterotrophs and subsequent thaumarchaeal oxidation of the resulting NH_4_^+^.

## RESULTS

### GROWTH EXPERIMENTS

Growth experiments were conducted with axenic cultures of *Nitrosopumilus* strains D3C and NF5 (Bayer *et al*., 2019a) grown in NH_4_^+^-free Synthetic Crenarchaeota Medium (SCM; Könneke *et al*., 2005) amended with a variety of single DON compounds (1 or 2 mM added N; Table 1) as their sole energy and N source. Small amounts of NO_2_^−^ (10-50 µM) were linearly produced over 70 days in all treatments (including negative controls with no added NH_4_^+^ or DON), presumably due to NH_4_^+^ contamination or breakdown and oxidation of media components such an antibiotics. Both strains converted NH_3_ into NO_2_^−^ stoichiometrically within 7 days, and D3C converted urea-N into NO_2_^−^ within ∼14 days (Table 1; Fig. S1), consistent with previous data from these isolates (Bayer *et al*., 2016; Bayer *et al*., 2019a). A small fraction (20-30%) of the glutamine-N amendment in both strains was converted to NO_2_^−^ linearly with time. None of the other DON amendments led to NO_2_^−^ production (Table 1; Fig. S1).

**Table 1.**
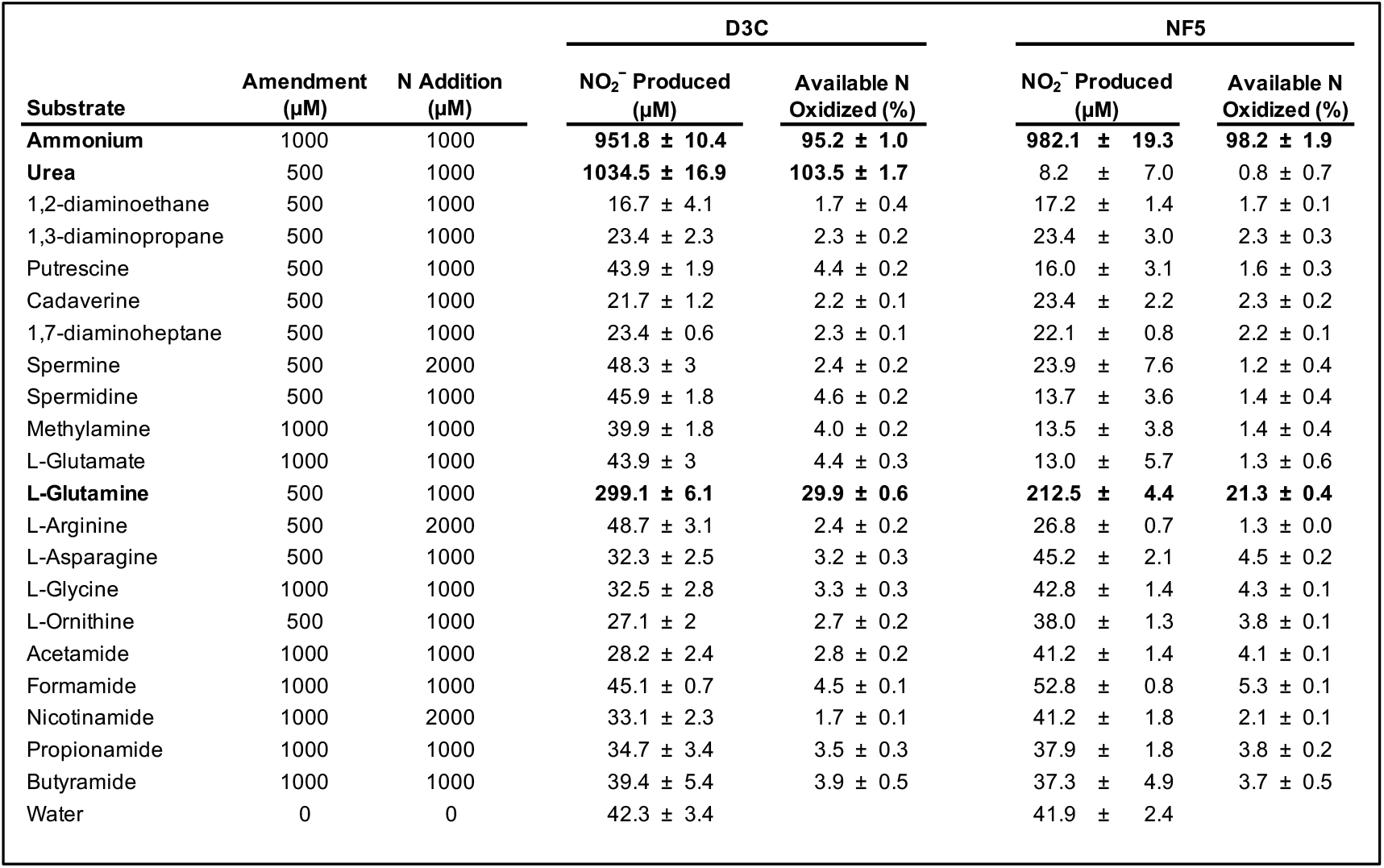
NO_2_ ^−^ produced (mean ± standard deviation of triplicates) by axenic thaumarchaeal strains in media containing single organic N compounds as the sole energy source. Bolded text shows treatments with at least one isolate accumulating NO_2_^−^ to a level greater than the negative control treatments.

Further growth experiments were conducted with enrichment cultures of *Nitrosopumilus* strains D3C and NF5 containing ∼5-15% bacterial 16S rRNA genes (Table 2) belonging to the heterotrophic bacterium *Oceanicaulis alexandrii* (Bayer *et al*., 2016). Growth on NH_3_ or urea proceeded rapidly and matched published rates (Bayer *et al*., 2016). NO_2_^−^ accumulation during growth of enrichments on NH_3_-free SCM amended with either 500 µM PUT or 500 µm PUT + 50 µM NH_4_^+^ was often linear and relatively slow (Fig. 1), but occasionally resembled a typical microbial growth curve (e.g., *N. adriaticus* NF5 grown with PUT and NH_4_^+^ amendment; Fig. 1 A). During growth on NH_3_, thaumarchaeal and bacterial 16S rRNA genes in selected treatments (those shown in Fig. 1 A, C) generally retained the same relative abundance as the respective inoculum (∼90-95% thaumarchaeal genes) and attained abundances comparable to the initial inoculum by the end of the experiment (Table 2). Thaumarchaeal and bacterial genes gradually increased in treatments containing PUT, with bacterial genes often reaching higher abundances than the initial inoculum (Table 2). Nitrite concentration increased rapidly when *N. adriaticus* NF5 was amended with PUT + NH_4_^+^ (Fig. 1 A). Both populations of genes increased in this treatment; notably, bacterial genes increased from 6.7×10^6^ genes mL^−1^ in the inoculum (8.2% of 16S genes) to 1.4– 4.5×10^8^ genes mL^−1^ (79.3–81.0% of 16S genes) during the incubation (Table 2).

**Table 2.** Quantification of bacterial and thaumarchaeal 16S rRNA genes via qPCR for selected experiments with enrichment cultures. NO_2_^−^ accumulation in these experiments is shown in Fig. 1 A, C.

| Strain | Substrate | Nutrients | Time (d) | 16S rRNA genes mL<sup>-1</sup> |  |  |
| --- | --- | --- | --- | --- | --- | --- |
|  |  |  |  | Bacterial | Thaumarchaeal | % Thaumarchaeota |
| D3C | All D3C treatments | Fig. 1 A | Inoculum | 1.5 × 10<sup>7</sup> | 9.4 × 10<sup>7</sup> | 86.3 |
| NF5 | All NF5 treatments | Fig. 1 A | Inoculum | 6.7 × 10<sup>6</sup> | 7.5 × 10<sup>7</sup> | 91.8 |
| NF5 | NH<sub>4</sub><sup>+</sup> | Fig. 1 A | 8 | 7.8 × 10<sup>6</sup> | 5.8 × 10<sup>7</sup> | 88.1 |
| NF5 | NH<sub>4</sub><sup>+</sup> | Fig. 1 A | 8 | 6.6 × 10<sup>6</sup> | 6.5 × 10<sup>7</sup> | 90.9 |
| NF5 | Putrescine + NH<sub>4</sub><sup>+</sup> | Fig. 1 A | 37 | 4.5 × 10<sup>8</sup> | 1.2 × 10<sup>8</sup> | 20.7 |
| NF5 | Putrescine + NH<sub>4</sub><sup>+</sup> | Fig. 1 A | 37 | 1.4 × 10<sup>8</sup> | 3.4 × 10<sup>7</sup> | 19.0 |
| D3C | All D3C treatments | Fig. 1 C | Inoculum | 1.5 × 10<sup>6</sup> | 1.3 × 10<sup>7</sup> | 94.2 |
| D3C | NH<sub>4</sub><sup>+</sup> | Fig. 1 C | 8 | 8.7 × 10<sup>5</sup> | 1.4 × 10<sup>7</sup> | 96.6 |
| D3C | NH<sub>4</sub><sup>+</sup> | Fig. 1 C | 8 | 1.3 × 10<sup>6</sup> | 1.5 × 10<sup>7</sup> | 95.5 |
| D3C | Urea | Fig. 1 C | 8 | 1.3 × 10<sup>6</sup> | 7.3 × 10<sup>6</sup> | 91.9 |
| D3C | Urea | Fig. 1 C | 8 | 1.7 × 10<sup>6</sup> | 7.9 × 10<sup>6</sup> | 90.1 |
| D3C | Urea | Fig. 1 C | 14 | 1.1 × 10<sup>6</sup> | 3.4 × 10<sup>7</sup> | 98.4 |
| D3C | Urea | Fig. 1 C | 14 | 1.2 × 10<sup>6</sup> | 3.7 × 10<sup>7</sup> | 98.4 |
| D3C | Urea | Fig. 1 C | 18 | 1.1 × 10<sup>6</sup> | 3.1 × 10<sup>7</sup> | 98.3 |
| D3C | Urea | Fig. 1 C | 18 | 1.3 × 10<sup>6</sup> | 4.0 × 10<sup>7</sup> | 98.3 |
| D3C | Putrescine | Fig. 1 C | 8 | 1.0 × 10<sup>6</sup> | 6.1 × 10<sup>6</sup> | 91.4 |
| D3C | Putrescine | Fig. 1 C | 8 | 1.8 × 10<sup>6</sup> | 8.9 × 10<sup>6</sup> | 90.9 |
| D3C | Putrescine | Fig. 1 C | 14 | 1.2 × 10<sup>6</sup> | 8.6 × 10<sup>6</sup> | 93.6 |
| D3C | Putrescine | Fig. 1 C | 14 | 1.1 × 10<sup>6</sup> | 8.3 × 10<sup>6</sup> | 93.6 |
| D3C | Putrescine | Fig. 1 C | 18 | 3.6 × 10<sup>6</sup> | 1.2 × 10<sup>7</sup> | 87.0 |
| D3C | Putrescine | Fig. 1 C | 18 | 1.4 × 10<sup>6</sup> | 1.4 × 10<sup>7</sup> | 95.0 |
| NF5 | All NF5 treatments | Fig. 1 C | Inoculum | 1.5 × 10<sup>6</sup> | 1.6 × 10<sup>7</sup> | 95.1 |
| NF5 | NH<sub>4</sub><sup>+</sup> | Fig. 1 C | 8 | 1.0 × 10<sup>6</sup> | 1.4 × 10<sup>7</sup> | 96.1 |
| NF5 | NH<sub>4</sub><sup>+</sup> | Fig. 1 C | 8 | 1.5 × 10<sup>6</sup> | 1.5 × 10<sup>7</sup> | 94.8 |
| NF5 | Putrescine | Fig. 1 C | 8 | 1.6 × 10<sup>6</sup> | 6.4 × 10<sup>6</sup> | 89.1 |
| NF5 | Putrescine | Fig. 1 C | 8 | 1.6 × 10<sup>6</sup> | 5.9 × 10<sup>6</sup> | 88.3 |
| NF5 | Putrescine | Fig. 1 C | 14 | 1.0 × 10<sup>6</sup> | 7.2 × 10<sup>6</sup> | 93.6 |
| NF5 | Putrescine | Fig. 1 C | 14 | 1.1 × 10<sup>6</sup> | 7.7 × 10<sup>6</sup> | 93.2 |
| NF5 | Putrescine | Fig. 1 C | 18 | 3.0 × 10<sup>6</sup> | 9.7 × 10<sup>6</sup> | 86.7 |
| NF5 | Putrescine | Fig. 1 C | 18 | 2.5 × 10<sup>6</sup> | 1.1 × 10<sup>7</sup> | 89.8 |

**Fig. 1.**
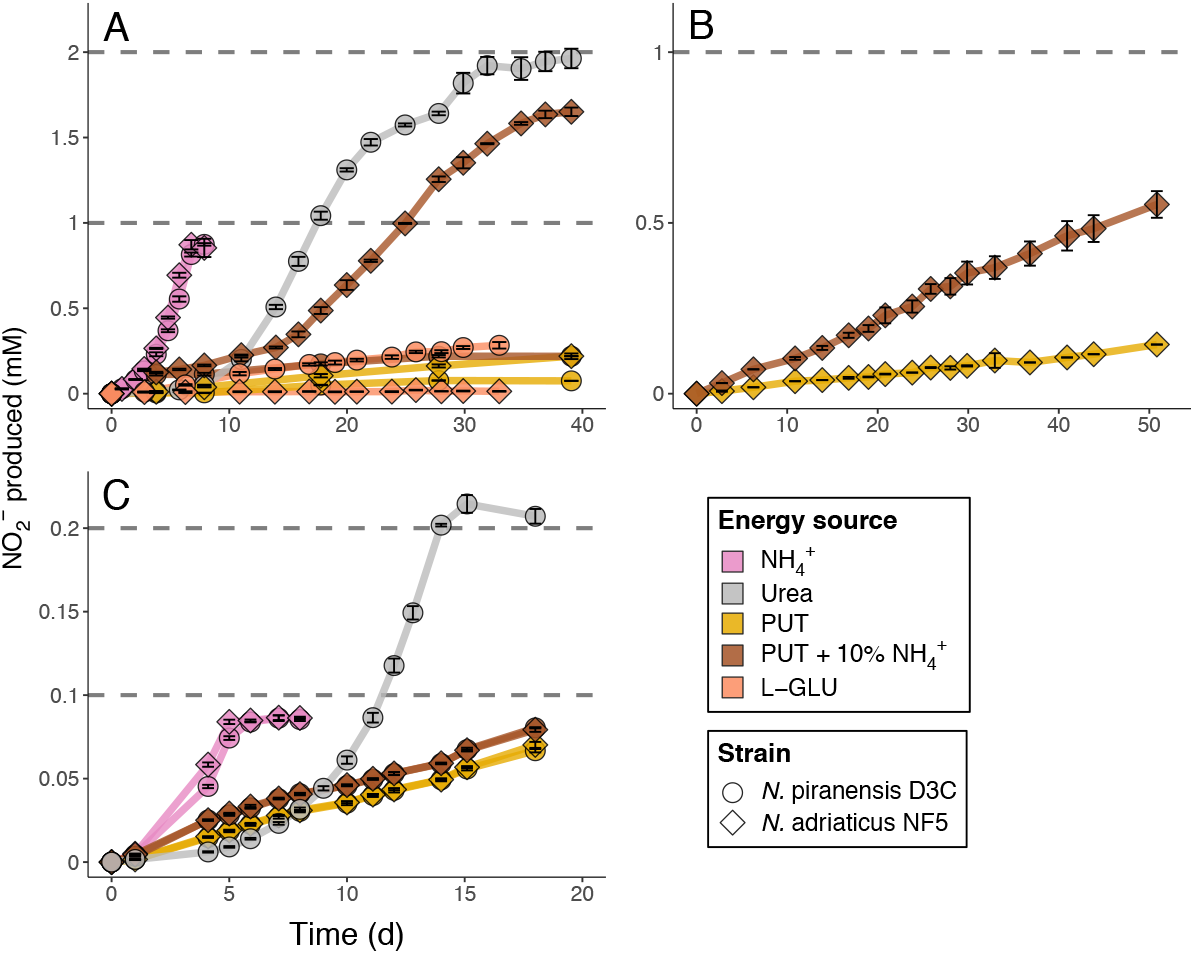
NO_2_ ^−^ accumulation by enrichment cultures of *N. piranensis* D3C (circles) and *N. adriaticus* NF5 (diamonds) over time. Color denotes the compound included as sole energy source in media. Points show the average of triplicate incubation bottles while error bars show standard deviation. Dashed lines show maximal NO_2_ ^−^ production given added N (two lines are present to show total added N when NH _4_^+^ and urea were included). **A)** Growth with either 1 mM NH_4_ ^+^, 1 mM urea, 1 mM PUT, 1 mM PUT plus 100 µM NH_4_ ^+^, or 1 mM L-GLU. **B)** Growth with either 1 mM PUT or 1 mM PUT plus 100 µM NH_4_ ^+^ (*N. adriaticus* NF5 only). **C)** Growth with either 100 µM NH_4_ ^+^, 100 µM urea, 100 µM PUT, or 100 µM PUT plus 10 µM NH_4_ ^+^. Bacterial and thaumarchaeal 16S rRNA gene quantities for selected experiments (panels A, C) are shown in Table 2.

### ISOTOPE EXPERIMENTS

Both axenic *Nitrosopumilus* strains oxidized 100 µM NH_3_ to NO_2_^−^ within 6 days. When 1.25 µM ^15^N-NH_4_^+^ was added to the growth media containing 100 µM unlabeled NH_4_^+^, all of the ^15^N was converted to ^15^N-NO_2_^−^ by stationary phase (NF5: 1.20 ^15^N-NO_2_^−^ µM produced; D3C: 1.09 µM; Fig. 2). Oxidation of polyamine-N was tested by adding ^15^N-labeled 1,2-diaminoethane (DAE), 1,3-diaminopropane (DAP), or PUT to the growth media. L-GLU was used as a control for remineralization, given its rapid catabolism and NH_4_^+^ remineralization by marine heterotrophs (Hollibaugh, 1978; Goldman *et al*., 1987; Goldman and Dennett, 1991). Growth with ^15^N-labeled DAE, DAP, or GLU did not produce ^15^N-NO_2_^−^, with values similar to the negative control (^14^N-NH_4_^+^ amendment, no added ^15^N). During growth with ^15^N-PUT, ∼35% of the added ^15^N was oxidized to ^15^N-NO_2_^−^ as NH_3_ was consumed (NF5: 0.45 ^15^N-NO_2_^−^ µM produced; D3C: 0.43 µM; Fig. 2).

**Fig. 2.**
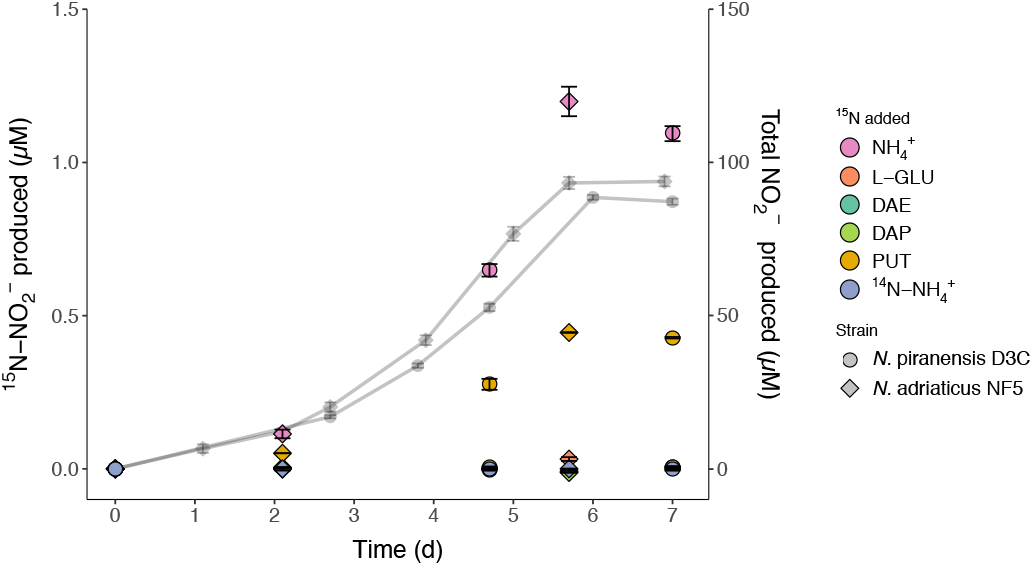
^15^N-NO_2_ ^−^ production from ^15^N-labeled organic N compounds during growth of axenic cultures of *N. piranensis* D3C (circles) and *N. adriaticus* NF5 (diamonds) on NH_3_. Gray points show total NO_2_^−^ (^14^ N and ^15^N; right y-axis) while points with color show concomitant production of ^15^N-NO_2_^−^ (left y-axis). All points represent the midpoint of duplicate incubation bottles, with error bars representing the range. The ^14^N-NH_4_^+^ treatment was a negative control containing no added ^15^N.

To test whether this ^15^N-PUT oxidation was due to abiotic reactions with short-lived reactive metabolic intermediates, ^15^N-PUT was incubated in filtered-sterilized, oligotrophic seawater or filtered-sterilized SCM dosed with nitric oxide (NO), peroxynitrite (ONOO^−^), or hydrogen peroxide (H_2_O_2_). Amendment with only NO_2_^−^ or PUT provided controls with no reactive compounds (Fig. 3 F). There was no significant difference in *δ*^15^N_NOx_ values between incubations of culture medium or filtered seawater receiving NO additions (*U*=26, p=0.574); therefore, NO addition experiments using either matrix were combined in data analyses.

**Fig. 3.**
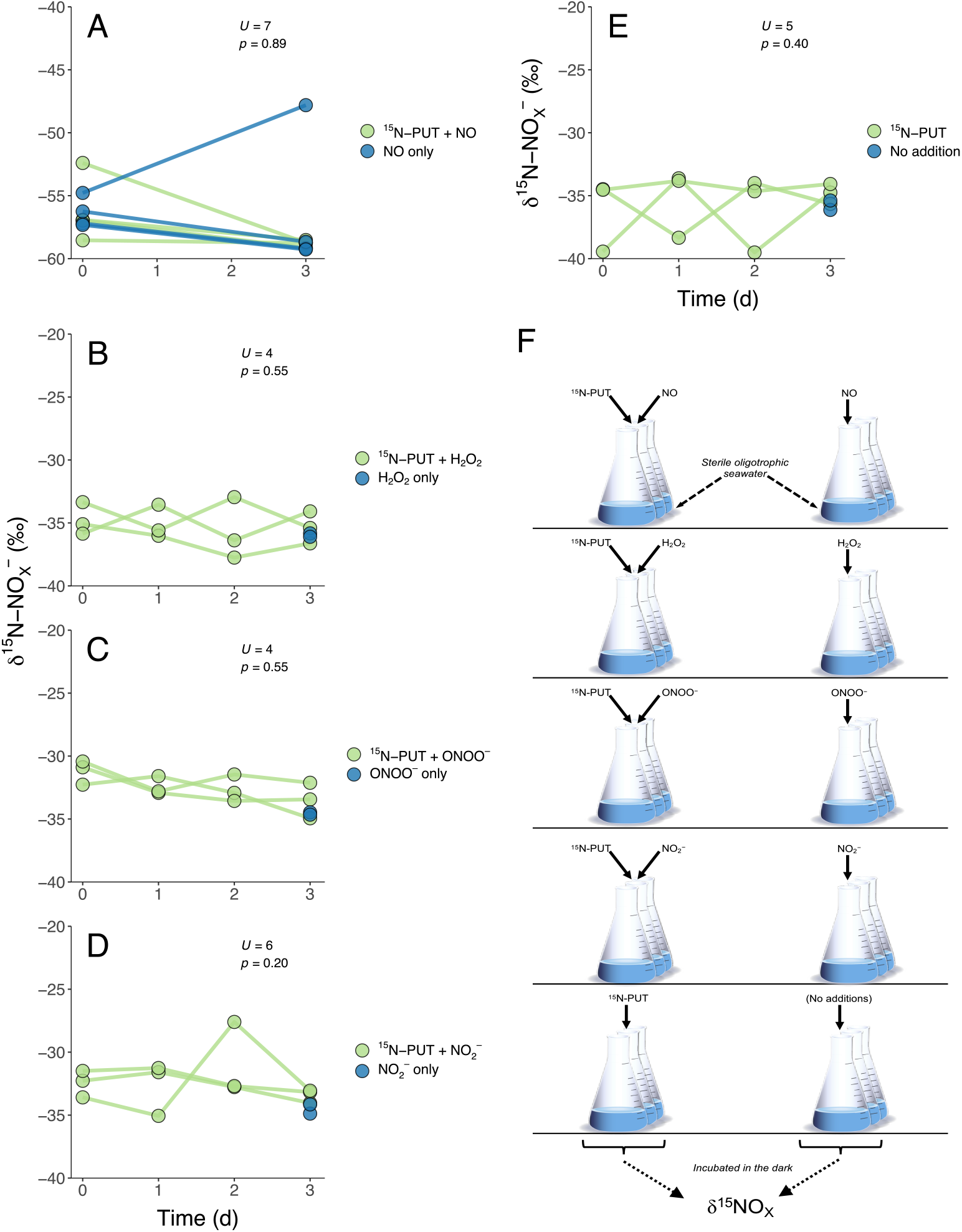
**A-E)** *δ*^15^NO_x_ (‰) over time in abiotic incubations containing ^15^N-labeled PUT and a single reactive compound (blue) or a reactive compound only (green). Results of a Mann-Whitney *U*-test are shown in each panel. **F)** Schematic of abiotic incubation experiments. The left column shows treatments including ^15^N-labeled PUT and a reactive compound while the right column shows control treatments containing a reactive compound and no ^15^N-labled PUT.

There was no significant difference in the ratio of ^15^N/^14^N in the nitrate plus nitrite (NO_X_) pool (*δ*^15^N_NOx_) at the termination of any of the incubations containing ^15^N-PUT versus those without ^15^N-PUT (Mann-Whitney *U*-tests, all *p*≥0.2; Fig. 3 A-E). Furthermore, there was no increase in *δ*^15^N_NOx_ values when NO gas was directly mixed with a variety of ^15^N-labeled polyamines, L-GLU, or NH_4_^+^ in sterile oligotrophic seawater (Mann-Whitney *U*-test, p=0.64; Fig. 4). NO_2_^−^ in NO-amended incubation endpoints was likely produced via spontaneous NO oxidation (Lewis and Deen, 1994), as NO_2_^−^ was not detectable in incubations without added NO (regardless of ^15^N addition). Therefore, NO_2_^−^ concentration in incubation endpoints likely reflects the concentration of the initial NO amendment, and was fairly well correlated to *δ*^15^N_NOx_ values (Spearman’s *ρ* = –0.70, *p*=0.02; Fig. 4).

**Fig. 4.**
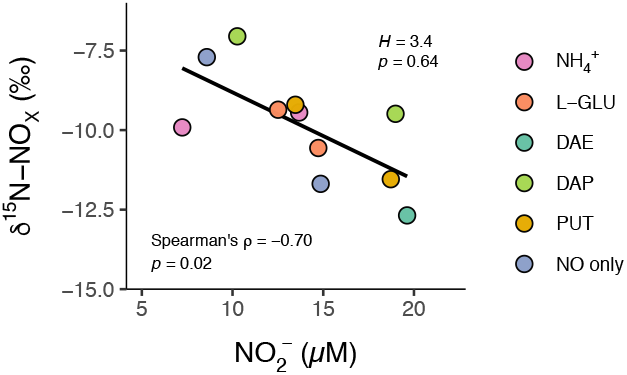
*δ*^15^NO_x_ (‰) as a function of NO_2_^−^ concentration following 24 hours of abiotic incubation of sterile seawater amended with ^15^N-labeled compounds and NO gas. NO (1% in N_2_) was directly bubbled into seawater. Color denotes ^15^N addition. Results of a Kruskal-Wallis test (comparing *δ*^15^NO_x_ between treatment groups) and Spearman’s correlation coefficient are shown. NO_2_^−^ in these incubation endpoints was likely due to abiotic NO oxidation and therefore reflects the amount of added NO.

## DISCUSSION

### *Organic N oxidation by* Thaumarchaeota *relies on remineralization*

Polyamines are ubiquitous in cells (Tabor and Tabor, 1985; Nishibori and Nishijima, 2004; Liu *et al*., 2016) and highly labile in aqueous environments (Lu *et al*., 2015; Liu *et al*., 2015; Krempaska *et al*., 2017; Madhuri *et al*., 2019). The amine groups at the ends of polyamines are essentially NH_3_ with an aliphatic C chain substituted for a hydrogen, and since their pK_a_ values (∼10.5, depending on chain length and temperature) are close to that of NH_3_ (∼9.25), they are similarly protonated in seawater. Thus, recent field evidence of rapid oxidation of polyamine-N in *Thaumarchaeota*-rich ocean waters (Damashek *et al*., 2019a) begs the question of whether these archaea can use polyamine-N directly as an energy source.

We tested this hypothesis using two *Nitrosopumilus* strains phylogenetically similar to the *Thaumarchaeota* typical of South Atlantic Bight inshore waters (Hollibaugh *et al*., 2011; Hollibaugh *et al*., 2014; Liu *et al*., 2018; Damashek *et al*., 2019b), where previously-measured polyamine-N oxidation rates were high (Damashek *et al*., 2019a). Efforts to grow *N. piranensis* D3C and *N. adriaticus* NF5 with single polyamines as their sole energy source conclusively demonstrated that they cannot grow by oxidizing N from PUT or other common polyamines (Fig. S1). Previous experiments with the same strains found only minimal [^3^H]leucine incorporation, particularly when archaea were starved of NH_3_, similarly suggesting no ability to use leucine as a carbon or energy source (Bayer *et al*., 2019a). There is long-standing biogeochemical and genomic evidence that some marine *Thaumarchaeota* may use some organic compounds as C sources (e.g., Ouverney and Fuhrman, 2000; Teira *et al*., 2004; Swan *et al*., 2014; Seyler *et al*., 2018; Dekas *et al*., 2019), and potentially even as an energy source (in addition to NH_3_; Dekas *et al*., 2019). As with polyamines, we found no evidence of growth by either *Nitrosopumilus* strain on a wide variety of common amino acids, amines, or amides (Table 1; Fig. S1). This suggests that NH_3_, urea, and cyanate remain the only known growth substrates of marine *Thaumarchaeota*.

The lack of evidence for direct oxidation of DON-N by pure cultures suggests that observed rates of DON-N oxidation (and potentially assimilation) by *Thaumarchaeota* in mixed communities are not a consequence of direct oxidation by *Thaumarchaeota* alone, but instead couple heterotrophic DON-N remineralization to NH_4_^+^ with subsequent thaumarchaeal oxidation of this newly-produced NH_4_^+^. Strong evidence for this conclusion comes from growth experiments with enrichment cultures: the presence of a heterotrophic bacterium led to a substantial accumulation of NO_2_^−^ in the medium and an increase in the abundance of bacterial 16S rRNA genes, especially when cultures were amended with both PUT and NH_4_^+^ (Fig. 1; Table 2). In contrast, pure cultures of the same *Thaumarchaeota* were unable to oxidize N supplied as PUT in the absence of NH_3_ (Fig. S1). In addition to indicating a reliance of thaumarchaeal DON-N oxidation on external remineralization, these findings may have similar implications for experiments measuring DON assimilation by *Thaumarchaeota* in mixed communities.

Polyamines are highly labile and energy-rich, as their backbone contains multiple fully-reduced C atoms. It is not surprising that adding this rich C source led to rapid bacterial growth in enrichment cultures, similar to the stimulation of heterotrophic communities documented in field studies (Mou *et al*., 2011; Mou *et al*., 2015). Notably, the C:N ratio of PUT (2) is lower than that of natural marine microbial cells (typically ∼4-5; Goldman *et al*., 1987; White *et al*., 2019), suggesting PUT catabolism by marine bacteria will lead to production of excess NH_4_^+^ (Hollibaugh, 1978; Goldman and Dennett, 1991). Therefore, NO_2_^−^ production in enrichment cultures could result from bacterial catabolism of PUT, followed by oxidation of the resulting NH_4_^+^ by *Thaumarchaeota*, typically leading to slow and linear NO_2_^−^ accumulation (Fig. 1). The occasional rapid accumulation of NO_2_^−^ when NH_4_^+^ was added alongside PUT (Fig. 1 A) may indicate the bacterium in this enrichment was unable to use PUT alone as an N source (i.e., needed free NH_4_^+^ to catabolize PUT), or may suggest bacterial dependence on *Thaumarchaeota* (growing on the low concentration of NH_3_) as a source of nutrients or organic substrates (e.g., Doxey *et al*., 2015; Heal *et al*., 2018; Bayer *et al*., 2019b).

Since many DON compounds are highly labile C sources, it is imperative to systematically account for rapid heterotrophic remineralization of N from DON in incubations of mixed communities before concluding that *Thaumarchaeota* are capable of direct oxidation or assimilation of DON, ideally by using pure cultures to test field hypotheses. Urea is a clear exception, since pure cultures of some marine *Thaumarchaeota* can grow with urea-N as their only energy and N source (Qin *et al*., 2017a; Bayer *et al*., 2019a). Direct cyanate-N oxidation/assimilation may also occur in marine *Thaumarchaeota*, but it is unclear how marine *Thaumarchaeota* could directly oxidize cyanate-N. While pure cultures of *Nitrosopumilus maritimus* SCM1 produced ^15^N-NO_2_^−^ when ^15^N-cyanate was added to their growth media, this only happened when they were already growing on NH_3_ (Kitzinger *et al*., 2019; see discussion below). The pathway of cyanate oxidation (or NH_4_^+^ production from cyanate) is unclear since their genomes lack canonical cyanase genes (Kitzinger *et al*., 2019).

Of the myriad polyamines, amino acids, amines, and amides we tested as thaumarchaeal growth substrates, the only compound leading to appreciable NO_2_^−^ accumulation was glutamine: when both axenic strains were supplied with glutamine as their sole energy source, there was a slow, linear accumulation of NO_2_^−^ (Fig. S1). Since the increase in NO_2_^−^ concentration over time did not resemble a growth curve, it is unlikely that the archaea were directly using glutamine-N as a growth substrate. Although the mechanism is not clear, glutamine was probably slowly remineralized to NH_4_^+^, which was then oxidized. A simple explanation is abiotic glutamine deamination, which occurs relatively rapidly in liquid media containing phosphate and bicarbonate at neutral pH (Gilbert *et al*., 1949), such as the SCM used here. However, our data cannot rule out a biological explanation. Glutamine is a common biochemical amino/amido donor, and *Thaumarchaeota* encode many genes involved in these reactions (e.g., glutamine amidotransferases or transaminases in amino acid and cofactor biosynthesis pathways; Walker *et al*., 2010; Kerou *et al*., 2016) as well as a variety of putative amino acid transporters (Offre *et al*., 2014). If excess glutamine is transported into thaumarchaeal cells, some may be recycled to NH_4_^+^ for either assimilatory or dissimilatory use. This amino acid “recycling” is common in energy-starved microbes (Lever *et al*., 2015) and was recently hypothesized to play a role in thaumarchaeal survival in deep sea sediments (Kerou *et al*., 2021). It is possible that energy-starved *Thaumarchaeota* in our experiment transported glutamine into their cells but were unable to incorporate it into biomass, so they hydrolyzed the glutamine amide N and shunted the resulting NH_4_^+^ into energy production, leading to slow growth. Direct tests of thaumarchaeal glutamine uptake and transformation compared to abiotic breakdown in SCM would be needed to uncover the mechanism behind the slow glutamine oxidation documented here.

### Potential co-metabolism or abiotic breakdown of polyamines

Although axenic cultures of marine *Nitrosopumilus* were unable to grow using PUT as their sole energy and N source, the significant fraction of ^15^N-PUT oxidized during growth on NH_3_ (Fig. 2), combined with the correlation between PUT-N and NH_3_ oxidation rates in the ocean (Damashek *et al*. 2019a), suggests PUT may be co-metabolized by the archaeal ammonia monooxygenase enzyme (AMO) or decomposed by reactive intermediates produced during NH_3_ oxidation (e.g., Martens-Habbena *et al*., 2015; Kim *et al*., 2016). Therefore, *Thaumarchaeota* in environments with high polyamine availability and enough NH_3_ for rapid growth may indirectly oxidize a significant amount of PUT-N to NO_2_^−^ despite their inability to use PUT-N as a direct energy source, increasing the flux of N from DON into the NO_X_ pool and potentially contributing to measured PUT-N oxidation rates in the ocean (Damashek *et al*., 2019a).

Co-metabolism of a variety of compounds due to non-specific oxidation by ammonia monooxygenase is well documented in ammonia-oxidizing bacteria (AOB). Some AOB can, for example, co-metabolize methane (Hyman and Wood, 1983; Ward, 1987), and some AOB and *Thaumarchaeota* can co-metabolize a variety of organic compounds (e.g., Rasche *et al*., 1991; Wright *et al*., 2020). Furthermore, co-metabolism of some DON compounds has been documented: two AOB strains and the thaumarchaeote *Nitrososphaera gargensis* can co-metabolize the tertiary amines mianserin and ranitidine during growth on NH_3_ (Men *et al*., 2016). Given the close structural and chemical similarity between NH_3_ and the primary amine groups on PUT-N, it seems conceivable that archaeal ammonia monooxygenase may similarly be able to oxidize PUT-N.

There has been recent recognition of the role played by reactive metabolic intermediates (primarily NO and H_2_O_2_) in thaumarchaeal physiology, and of the importance of chemical transformations catalyzed by these reactive compounds in culture experiments. For example, pure thaumarchaeal isolates produce H_2_O_2_ (Kim *et al*., 2016; Bayer *et al*., 2019c) but lack catalase to facilitate its rapid detoxification. Exposure of isolates or field populations to high concentrations of H_2_O_2_ therefore arrests their growth and activity (Kim *et al*., 2016; Tolar *et al*., 2016; Qin *et al*., 2017b; Bayer *et al*., 2019c). NO, an obligate intermediate in the archaeal ammonia oxidation pathway, accumulates during growth of thaumarchaeal cultures (Martens-Habbena *et al*., 2015; Kozlowski *et al*., 2016; Sauder *et al*., 2016; Hink *et al*., 2017) and is found in high concentrations in marine regions with high nitrification rates or abundant *Thaumarchaeota* (Ward and Zafiriou, 1988; Lutterbeck *et al*., 2018). NO is a reactive radical that rapidly forms reactive nitrogen oxide species (RNOS; e.g., NO_2_ or N_2_O_3_) upon exposure to O_2_ or superoxide (Wink and Mitchell, 1998). RNOS are highly destructive of many biological molecules; relevant to our data, RNOS react with primary amines to produce nitrosamines that are subsequently deaminated to NH_4_^+^ (Ridnour *et al*., 2004). These reactions may be a mechanism for liberating NH_4_^+^ from polyamines due to NO production during thaumarchaeal growth on NH_3_. Similarly, reactions of NO with various compounds have been posited to explain chemical transformations during thaumarchaeal growth: the abiotic reaction between NO and reduced iron in growth media is thought to form nitrous oxide in cultures (Kozlowski *et al*., 2016), and NO may react with cobalamin within thaumarchaeal cells, leading to nitrocobalamin production (Heal *et al*., 2018).

Given the capability of NO-derived RNOS to react with primary amines, speculation about the reactivity of NO in thaumarchaeal cultures, and the documented ability of polyamines to scavenge oxygen radicals (Khan *et al*., 1992; Ha *et al*., 1998; Chattopadhyay *et al*., 2003), we hypothesized that NO produced during NH_3_ oxidation could react abiotically with ^15^N-PUT to produce the ^15^N-NO_2_^−^ we observed (Fig. 2). However, experiments with cell-free artificial media and sterilized seawater showed no evidence of reaction between NO and ^15^N-polyamines (Fig. 3 A, Fig. 4). NO_2_^−^ was produced rapidly in treatments with added NO since NO auto-oxidizes to NO_2_^−^ and H^+^ in oxic water (Lewis and Deen, 1994), but this NO_2_^−^ did not contain the ^15^N initially added as ^15^N-PUT. The correlation between NO_2_^−^ and *δ*^15^N_NOx_ values shown in Fig. 4 thus reflects the *δ*^15^N value of spontaneously oxidized NO, with no apparent ^15^N enrichment due to oxidation of any ^15^N-labeled substrates. We then tested whether ^15^N-polyamines react with H_2_O_2_ or ONOO^−^ (formed by reacting NO_2_^−^ and H_2_O_2_; Robinson and Beckman, 2005) to yield ^15^N-NO_2_^−^, as these reactive compounds are also produced during thaumarchaeal growth (Kim *et al*., 2016; Heal *et al*. 2018; Bayer *et al*., 2019b), but found no detectable ^15^N-NO_2_^−^ in these treatments either (Fig. 3). This suggests that some PUT oxidation seen in field experiments (Damashek *et al*. 2019a), with mixed cultures (Fig. 1), or during growth on NH_3_ (Fig. 2) may be explained by enzymatic co-metabolism of PUT, or due to reactions with reactive species that we did not test (e.g., superoxide or other RNOS compounds).

Our isotope experiments were restricted to PUT, DAP, and DAE due to limited commercial availability of ^15^N-labeled polyamines. Of these, only PUT is commonly found in high concentrations in phytoplankton and bacterial cells or marine waters (Nishibori *et al*., 2001; Lu *et al*., 2014; Liu *et al*., 2016; Lin and Lin, 2019). It remains unknown whether other common polyamines (e.g., spermine, spermidine, norspermine, or norspermidine) are comparably oxidized during thaumarchaeal NH_3_ oxidation. However, the oxidation of cyanate-N by marine *Thaumarchaeota* may be analogous, given that axenic *N. maritimus* SCM1 cultures only oxidized ^15^N-cyanate while growing on NH_3_ (Kitzinger *et al*., 2019), comparable to our results with ^15^N-PUT. Kitzinger *et al*. (2019) hypothesized that *N. maritimus* could break down cyanate extracellularly, since ^15^N-NH_4_^+^ was produced in cultures amended with ^15^N-cyanate despite no known cyanate hydratase genes existing in the *N. maritimus* SCM1 genome. Whether due to abiotic reactions with metabolic intermediates (intra- or extracellularly), some as yet undiscovered mechanism of extracellular remineralization, or co-metabolism by ammonia monooxygenase, the dual evidence of oxidation of PUT-N (Fig. 2) and cyanate (Kitzinger *et al*., 2019) during thaumarchaeal growth on NH_3_ suggests some DON compounds can be oxidized indirectly by growing *Thaumarchaeota*, but definitive demonstrations of mechanisms remain elusive.

## CONCLUSIONS

Given the ability of *Thaumarchaeota* to grow using reduced N supplied as urea and cyanate, there has been interest in their potential to access other forms of organic N, spurred by numerous field studies reporting putative thaumarchaeal DON assimilation or oxidation. Our experiments with two axenic isolates suggest marine *Thaumarchaeota* cannot directly access N supplied as polyamines, amino acids, amides, or primary amines as an energy source. Inclusion of a heterotrophic bacterium in enrichment cultures of *Thaumarchaeota* resulted in the oxidation of PUT-N, demonstrating the importance of DON remineralization to NH_4_^+^ by heterotrophs for *Thaumarchaeota* to oxidize N supplied as DON. Therefore, claims of DON use by *Thaumarchaeota* in mixed communities must strictly account for heterotrophic remineralization. Despite lacking the ability to grow on PUT alone, the surprising finding that both pure strains oxidized ^15^N supplied as PUT while growing on NH_3_ suggests some DON may be co-metabolized or broken down abiotically, possibly mediated by reactive species produced during NH_3_ oxidation. Abiotic experiments ruled out some of the known reactive oxygen and nitrogen intermediates of thaumarchaeal metabolism as oxidants, but did not identify the mechanism leading to ^15^N-PUT oxidation in cultures. This study suggests that oxidation of most DON-N for energy conservation by marine *Thaumarchaeota* requires initial DON remineralization to NH_4_^+^ by heterotrophs, but also indicates a potential role for co-metabolism or reactive metabolic byproducts in thaumarchaeal DON-N oxidation.

## EXPERIMENTAL PROCEDURES

### GROWTH EXPERIMENTS

Axenic cultures of *Nitrosopumilus piranensis* D3C and *N. adriaticus* NF5 (Bayer *et al*., 2019a) were grown in SCM amended with an array of single DON compounds as sole energy and N sources, including NH_4_^+^, urea, amino acids, primary amines (including polyamines), or amides added to 1 or 0.5 mM final concentration (1 or 2 mM N; Table 1). All treatments were run in triplicate for each strain and included pyruvate (200 µM) to scavenge reactive oxygen species (Kim *et al*., 2016) and 50 µg/mL each of streptomycin and kanamycin. NH_4_Cl was added to positive controls but was not included in other treatments. Experiments were initiated using a 10% (v/v) transfer of stationary phase cultures (with NH_4_^+^ completely consumed) to ensure against NH_4_^+^ carryover. Growth experiments were conducted in the dark at 29°C for ∼70 days and subsampled for NO_2_^−^ determination as described above. Culture purity was tested throughout the duration of the experiment by adding Marine Broth 2216 to aliquots of the culture (10% v/v) and monitoring for bacterial growth by measuring OD_600_.

Growth experiments were also conducted with early enrichment cultures of *N. piranensis* strain D3C and *N. adriaticus* strain NF5, in which approximately 5-15% of the cells were the heterotrophic bacterium *Oceanicaulis alexandrii* (Bayer *et al*., 2016; Bayer *et al*., 2019c). Triplicate cultures were grown in SCM containing either 1 mM NH_4_^+^, 1 mM urea, 1 mM PUT, and 1 mM PUT + 100 µM NH_4_^+^ as N or energy sources. Experiments were conducted at 29°C in the dark and subsampled over time for immediate NO_2_^−^ determination using standard methods (Griess reagent; Strickland and Parsons, 1972). In selected rounds of these experiments, subsamples were taken for qPCR analysis by mixing 0.8 mL of a culture with 0.8 mL 2X lysis buffer (1.5 M sucrose, 80 mM EDTA, 100 mM Tris; pH 8.3) and freezing immediately at −80°C. DNA was extracted using standard phenol/chloroform techniques (Tolar *et al*., 2013; Damashek *et al*., 2019a). Bacterial and thaumarchaeal 16S rRNA genes were quantified using primers BACT1369F/PROK1492R/TM1389F and GI_334F/GI_554R/TM519AR (Suzuki *et al*., 2000), respectively (see Table S1 for amplification conditions). Reactions were run in triplicate on a C1000 Touch Thermal Cycler/CFX96 Real-Time System (Bio-Rad) using the Platinum qPCR SuperMix-UDG (Thermo Fisher). Standard curves consisted of a dilution series of a linearized plasmid containing a previously-sequenced amplicon.

### ISOTOPE EXPERIMENTS

The ability of *Thaumarchaeota* to oxidize DON during growth on NH_3_ was assessed using ^15^N isotope tracers (98-99% ^15^N, Cambridge Isotope Laboratories). Axenic cultures were grown on SCM as described above except using 100 µM ammonium chloride and 20 µM pyruvate. Culture purity was tested as described above. Triplicate bottles per substrate were amended to 1.25 µM of ^15^N-labeled substrate: DAE, DAP, PUT, ammonium chloride, L-GLU, or ^14^N ammonium chloride (a negative control with no added ^15^N). Both amine groups of DAE, DAP, and PUT were ^15^N-labeled, leading to total ^15^N additions of 2.5 µM in these treatments. Growth was estimated by measuring NO_2_^−^ production over time. At multiple timepoints, subsamples were frozen at −80°C in polypropylene tubes for isotopic analysis.

*δ*^15^NO_x_ values were measured using the bacterial denitrifier method (Sigman *et al*., 2001). Briefly, NO_X_ was converted to nitrous oxide by *Pseudomonas aureofaciens* and its N isotopic ratio was measured using a Finnigan MAT-252 isotope ratio mass spectrometer coupled with a modified GasBench II interface (Casciotti *et al*., 2002; Beman *et al*., 2011; McIlvin and Casciotti, 2011). The concentration of ^15^N-NO_2_^−^ (µM) was calculated by multiplying the NO_X_ atom fraction ^15^N by the NO_2_^−^ concentration (nitrate was not detectable).

Abiotic oxidation of ^15^N-PUT by multiple reactive compounds was tested by amending triplicate flasks containing 75 mL of 0.22-µm filtered aged surface water from the Gulf Stream or Station ALOHA (NH_4_^+^ and NO_X_ below the limit of detection), or sterile SCM, with ^15^N-PUT and a single reactive compound (NO, H_2_O_2_, or ONOO^−^; Fig. 3 F). Flasks were then incubated at 23°C in the dark. Two control treatments were run that did not contain reactive compounds: one amended with NO_2_^−^, and one with no addition. For each treatment (including the two negative controls), parallel replicated incubations were conducted with and without added ^15^N-PUT (Fig. 3 F). NO was generated from the NO donor (Z)-1-[N-(3-aminopropyl)-N-(n-propyl)amino]diazen-1-ium-1,2-diolate (PAPA NONOate; Cayman Chemical, Ann Arbor). 100 µM of PAPA NONOate was added to each flask to produce 200 µM NO (Hrabie *et al*., 1993). The PAPA NONOate stock was prepared by slowly injecting 5.7 mL 0.01 M NaOH into an airtight vial containing 50 mg PAPA NONOate. This stock was wrapped in foil and stored at 4°C prior to use within hours. NO addition experiments were also conducted using blank SCM media (no added N, as described above). In H_2_O_2_ (Millipore Sigma, Burlington, MA), ONOO^−^, NO_2_^−^, and no addition control treatments, 1 µM of the respective compound was added to flasks containing 100 nM ^15^N-PUT, while NO treatment flasks contained 1.25 µM ^15^N-PUT. ONOO^−^ was generated by mixing NO_2_^−^ and H_2_O_2_ (Robinson and Beckman, 2005). At the beginning and end of each incubation, subsamples were frozen at −80°C for *δ*^15^NO_x_ determination (as described above). Twenty µM sodium nitrate with a known *δ*^15^N_NOx_ value was added to samples prior to conversion of NO_X_ to nitrous oxide to enable isotopic measurements in experiments with no NO_2_^−^ accumulation (H_2_O_2_, ONOO^−^, NO_2_^−^, and no addition control).

In tests of the oxidative ability of NO on multiple ^15^N-labeled compounds, NO gas (1% v/v in N_2_; Airgas, Radnor Township, PA) was directly bubbled through a syringe into Erlenmeyer flasks containing filtered aged surface water from the Gulf Stream (NH_4_^+^ and NO_X_ below the limit of detection). ^15^N-labeled NH_4_^+^, L-GLU, DAE, DAP, and PUT were added to duplicate flasks (as well as a negative control with no added ^15^N) and incubated at 23°C in the dark. After 24 hours, subsamples were frozen at −80°C for *δ*^15^NO_x_ determination (as described above).

### DATA ANALYSIS

Mann-Whitney *U*-tests were conducted using R (R Core Team, 2019) to determine whether *δ*^15^NO_x_ values differed between abiotic incubation endpoints of treatments with and without ^15^N-PUT (shown in Fig. 3) and between abiotic incubation endpoints of treatments containing different ^15^N-labeled substrates (shown in Fig. 4). Correlations were determined by calculating Spearman’s rank correlation coefficient (*ρ*) in R. Plots were made using the ggplot2 R package (Wickham, 2016).

## ACKNOWLEDGMENTS

This work was funded by NSF OCE grants 1538677 (to JTH) and 1537995 (to BNP). BB was supported by the Uni:docs Fellowship of the University of Vienna. A contribution number from the University of Hawai’i School of Earth Science and Technology will be provided upon manuscript acceptance. The authors declare no conflicts of interest.

## SUPPLEMENTAL TABLES AND FIGURES

**Table S1**

Primers, cycling information, and assay data for qPCR reactions

**Fig. S1** NO_2_^−^ accumulation in axenic cultures of *N. piranensis* D3C (circles) and *N. adriaticus* NF5 (diamonds) over time. Colors denote the compounds added to the media as sole energy and N sources. Points show the average of triplicate incubation bottles while error bars show standard deviation.

