## Supplementary figures and images for "Limited accessibility of nitrogen supplied as amino acids, amides, and amines as energy sources for marine *Thaumarchaeota*"

### Supplemental Fig. S1

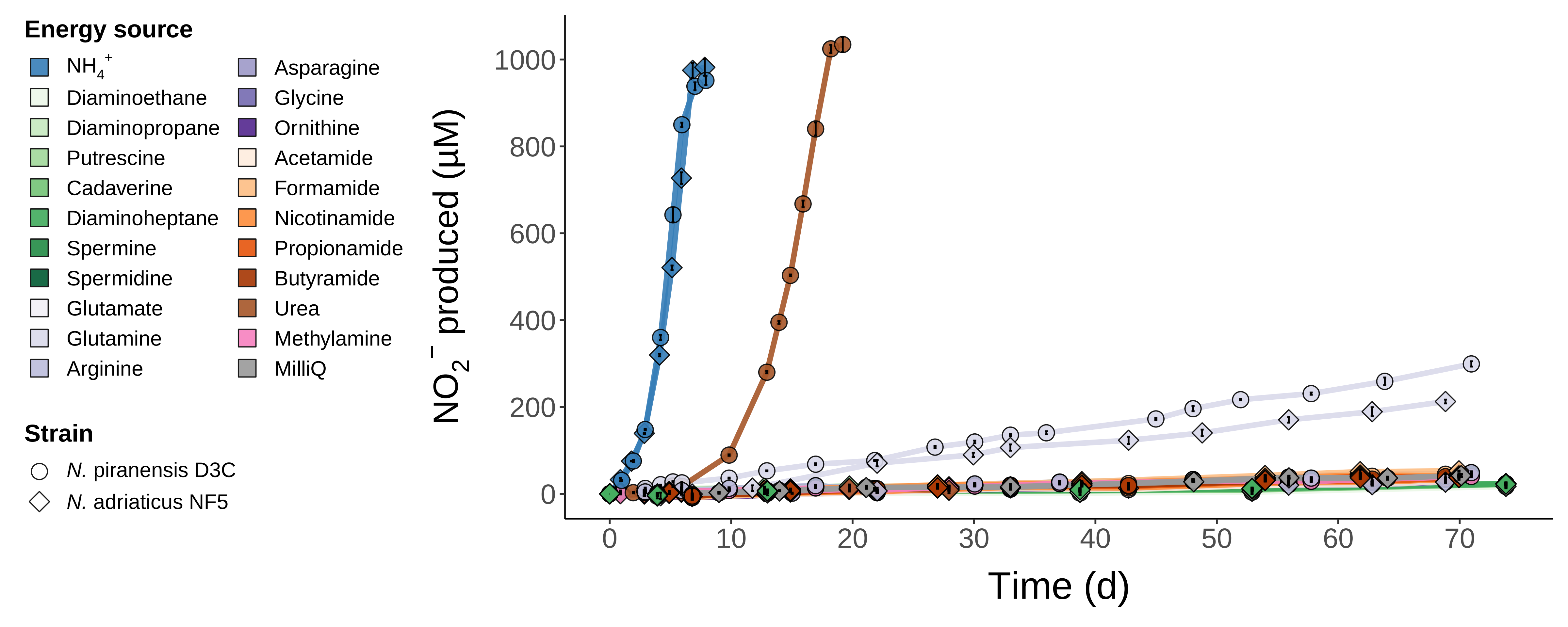
